# Effects of transcranial alternating current stimulation on spiking activity in computational models of single neocortical neurons

**DOI:** 10.1101/2021.01.21.427521

**Authors:** Harry Tran, Sina Shirinpour, Alexander Opitz

## Abstract

Neural oscillations are a key mechanism for information transfer in brain circuits. Rhythmic fluctuations of local field potentials control spike timing through cyclic membrane de- and hyperpolarization. Transcranial alternating current stimulation (tACS) is a non-invasive neuromodulation method which can directly interact with brain oscillatory activity by imposing an oscillating electric field on neurons. Despite its increasing use, the basic mechanisms of tACS are still not fully understood. Here, we investigate in a computational study the effects of tACS on morphologically realistic neurons with ongoing spiking activity. We characterize the membrane polarization as a function of electric field strength and subsequent effects on spiking activity in a set of 25 neurons from different neocortical layers. We find that tACS does not affect the firing rate of investigated neurons for electric field strengths applicable to human studies. However, we find that the applied electric fields entrain the spiking activity of large pyramidal neurons at < 1mV/mm field strengths. Our model results are in line with recent experimental studies and can provide a mechanistic framework to understand the effects of oscillating electric fields on single neuron activity. They highlight the importance of neuron morphology in responsiveness to electrical stimulation and suggest that large pyramidal neurons are most likely the prime target for tACS.

## Introduction

Brain oscillations are an important mechanism for brain communication. They arise from cyclical variations of local field potentials (LFPs) and are known to affect spiking activity by means of de- or hyperpolarization of neural membranes (Draguhn & Buzsáki, 2004). An increasing number of psychiatric diseases (e.g. schizophrenia) are being understood as deviations from healthy neural oscillatory activity (Uhlhaas & Singer, 2006). Thus, tools to directly interact with brain oscillations hold promise for new therapeutic approaches to harmonize pathological brain rhythms. Consequently, there has been a rising interest in non-invasive brain stimulation methods to modulate brain oscillations in humans. One popular method, transcranial alternating current stimulation (tACS), works by passing a weak oscillating electric current through electrodes attached to the scalp, creating an electric field inside the brain (Antal & Paulus, 2013). This electric field interacts with the spiking activity of neurons through transient membrane de- or hyperpolarization at the applied stimulation frequency.

Many experimental studies have investigated the effects of oscillating electric fields either *in-vitro* (Chan, Hounsgaard, 1988; Deans et al., 2007) or *in-vivo* animal studies (Fröhlich & McCormick, 2010; Johnson et al., 2020; Krause et al., 2019; Simal Ozen et al., 2010). One of the main hypotheses for tACS is the idea of neural entrainment. Entrainment means that neural activity at either the single neuron or population level synchronizes to the applied stimulation waveform (Chan & Nicholson, 1986; S. Ozen et al., 2010). However, entrainment is only one specific form of more general changes in spike-timing, meaning a deviation from regular spiking activity due to the applied stimulation. Thus, despite a growing body of experimental studies, the exact mechanisms involved in the alteration of neural activity due to tACS are still not fully understood.

Computational modeling is a key complementary tool to experimental work by providing a theoretical framework in which experimental findings can be understood. Modeling further allows to efficiently explore a larger range of stimulation parameters not feasible in experimental studies. Several computational studies have explored the effects of tACS on single neurons or neural populations (Ali et al., 2013; Cakan & Obermayer, 2019).These studies have highlighted the idea of network resonance in neural populations due to the applied oscillating electric field. Common to these studies is the use of simplified neuron models such as the ball-and-stick model (Aspart et al., 2016) or the two-compartment model (Ladenbauer & Obermayer, 2019). While such models are well suited for modelling large neuronal populations, they can give little insight about single neuron responses. This is due to the lack of a detailed dendritic morphology which strongly shapes neural activity (Ostojic et al., 2015). Further, differences in neural morphology and biophysics cannot be captured by simplified models but require detailed individualized neuron models. Thus, in order to study the effects of tACS on the single neuron level in different cell types, it is necessary to develop morphologically realistic neuron models with detailed biophysics.

Here, we conduct a computational study to investigate the effect of tACS on single-neuron spiking activity in a set of morphologically realistic neocortical neurons using cable theory to accurately model the underlying biophysics (Rall, 1962). Ongoing spiking activity was created through synaptic input randomly located in the dendritic branches. We explore the effect of electric field strength on neural activity at 10 Hz stimulation frequency and investigate neural entrainment and changes in spike timing and firing rate. Our computational study can thus give important insights into tACS mechanisms.

## Methods

Matlab code and NEURON code for neural activity modeling and analysis are freely available at GitHub: https://github.com/htran1902/tACS_effects_on_single_neurons.

### Neuron models and synaptic activity modeling

We used multi-compartmental conductance-based neuron models with realistic morphologies from all cortical layers generated in the NEURON environment (Carnevale & M.L. Hines, 1997). These models were adapted from ModelDB (Aberra et al., 2018, accession number 066023). In these neuron models, morphologies are based on the reconstructions from the Blue Brain Project (Markram et al., 2015). In summary, the database includes a set of 25 neocortical neurons composed of five neurons for each neocortical layer): Layer 1 neurogliaform cell (L1 NGC), Layer 2/3 pyramidal cell (L2/3 PC), Layer 4 large basket cell (L4 LBC), Layer 5 pyramidal cell (L5 PC) and Layer 6 pyramidal cell (L6 PC) (see supplementary figure 1 for the morphologies). Different ion channels and myelination are modelled as outlined in detail in (Aberra et al., 2018; Markram et al., 2015). Furthermore, we modified these models to incorporate oscillating external electric fields, and intrinsic synaptic activity. To this end, a synapse was added to each of the 25 cells at a random location on the apical dendrite for pyramidal cells or the basal dendrite for interneurons. However, to account for the various cell morphologies we varied the distance between the soma and the synaptic input to the following: (median distance from the soma for L1 NGC: 17.62 *μm*, L2/3 PC: 27.97 *μm*, L4 LBC: 35.08 *μm*, L5 PC: 46.62 *μm*, L6 PC: 13.69 *μm*). We modeled the synaptic activity within a single postsynaptic compartment on the cell body by using a two-exponential function as follows (Destexhe et al., 1998; Rothman, Jason S and Silver, 2014). The synaptic current that results from the chemical synapse is given by the following expressions:

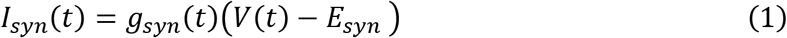

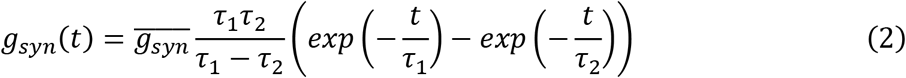

The parameter *E_syn_* corresponds to the synaptic reversal potential, *g_syn_*(*t*) is the time-dependent synaptic conductivity, 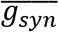 is the maximum synaptic conductance, *V*(*t*) is the membrane potential and τ_1_ and τ_2_ are respectively the rise time and the decay time. Here, we only used excitatory synapses and implemented them by setting *E_syn_* to 0mV, τ_1_ = 2ms and τ_2_ = 10*ms*. The synaptic weight was chosen according to the cell morphology and type in order to have an average firing rate between 5 and 13 spikes per second.

### Modeling of tACS

The frequency of the tACS oscillation was set to 10Hz. We simulated a range of electric field strengths between 0 mV/mm and 3 mV/mm. The amplitude was increased by 0.1 mV/mm steps between 0 mV/mm and 2 mV/mm, and 0.2 mV/mm steps between 2 mV/mm and 3mV/mm. We chose this range to cover intensities typically induced in human and animal studies as well as to explore a higher dosing range which might be achieved in future tACS applications. The electric field is spatially uniform and aligned with the y-axis which corresponds to the somatodendritic axis of the pyramidal cells (Figure 1A). The total duration of the stimulation was 8min and divided in two equal periods: a 4min tACS-free period and a 4min period with tACS (Figure 1.B). We used the extracellular mechanism of NEURON to simulate the interaction between the electric field and the cell activity (Anastassiou et al., 2010; Nagarajan et al., 1993). The NEURON environment is based on cable theory which discretizes the neuron morphology into small compartments on which the neural dynamics are computed. The membrane voltage at each compartment is approximated with the voltage at its geometrical center. The relation between the extracellular membrane voltage and the amplitude of the electric field is linear and is given by 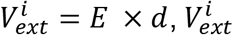 is the extracellular potential of the node *j*, *E* is the electric field strength and *d* is the distance between the cathode and the node *j* (Wang et al., 2018).

**Figure 1.**
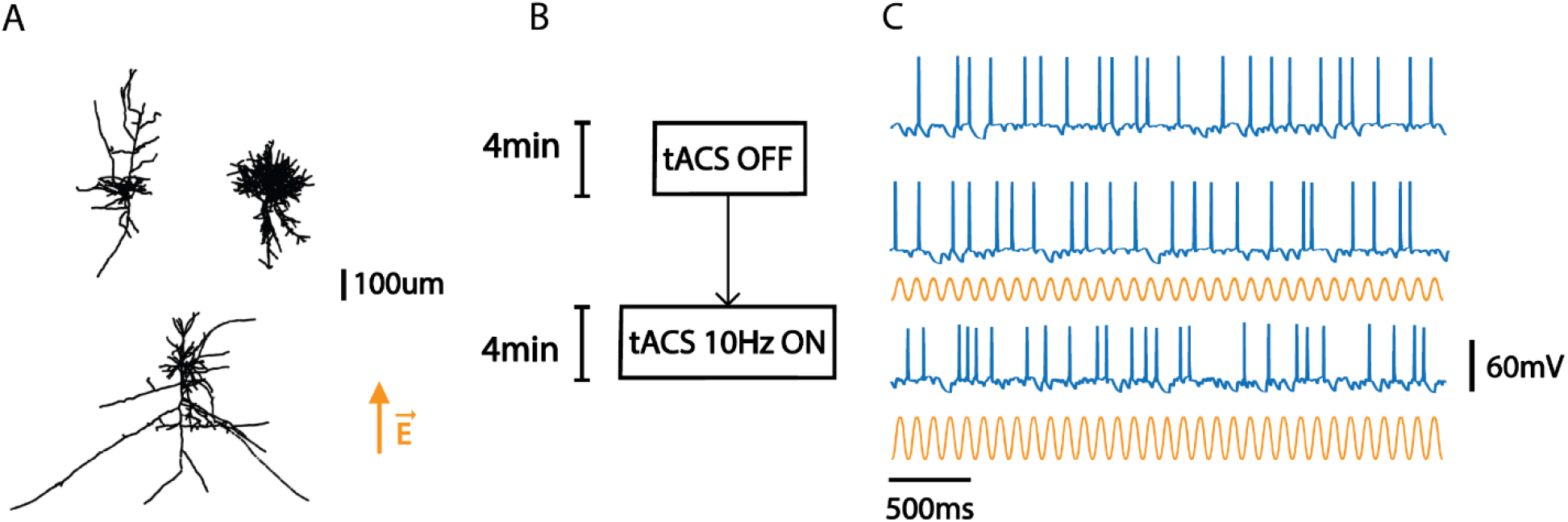
tACS simulation setup. **(A)** Examples of the complex realistic neocortical neuron morphologies used in the simulations. Top left. L6 pyramidal cell. Top right. L4 large basket cell. Bottom. L2/3 pyramidal cell. The orange arrow indicates the direction of the applied electric field along the y-axis. **(B)** A 10Hz oscillating electric field was applied to each neocortical cell for 4min. For comparison another 4min without tACS was simulated. **(C)** Spiking activity was generated through a randomly located synapse with a Poisson distribution. Top. Baseline neural spiking activity. Middle. Spiking activity for an electric field strength of 0.5 mV/mm. Bottom. Spiking activity for an electric field strength of 1 mV/mm.

### Characterizing the membrane polarization

In order to determine the membrane polarization with respect to the applied electric field strength, we used the same cells but without any synaptic input (silent neuron). For each cell, we recorded the somatic polarization for a given electric field strength. It has been shown in previous experimental studies that the somatic membrane potential increases linearly with the applied electric field strength (Bikson et al., 2004; Radman et al., 2009). This relationship is characterized through the polarization length *λ_p_* which measures the somatic polarization per unit electric field (unit: millimeters, mV *(mV/mm)^−1^). The higher *λ_p_*, the easier it is to depolarize the neuron. It has been shown that the polarization length can differ strongly across different cell types (Radman et al., 2009).

### Calculating phase-locking of spiking activity to the applied electric field

Multiple methods have been developed to measure neural oscillatory synchronization (Lowet et al., 2016; Picinbono, 1997). To quantify the neural entrainment, we computed the phase-locking value (PLV) which estimates spike timing relative to an oscillation, in our case the tACS waveform. The PLV is calculated as follows (Lowet et al., 2016)]:

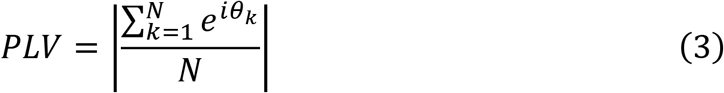

Where N is the number of action potentials and *θ_k_* is the phase of the tACS stimulation at which the k^th^ action potential occurs. A PLV of 0 means that no synchronization is existent – e.g. in case the spike times are uniformly distributed across the tACS cycle – while a value of 1 means perfect synchronization.

### Analysis of the neuron morphology

To identify morphological factors which can explain some of the observed differences in spiking activity due to tACS, we performed a quantitative analysis of the neuron morphology. For this we characterized the neuron morphologies through their effective lengths (units: *μm*) along the direction of the electric field (y-axis) as well as perpendicular (x-axis) to it. The effective lengths *L_X_* and *L_Y_* correspond to the distance between the two furthest points along the x-axis and y-axis, respectively. A high effective length means that a neuron has a morphology specifically oriented in that axis. We further calculated *ρ_YX_* as the ratio between *L_Y_* and *L_X_*. A high ratio means that the neuron has an elongated morphology in that a specific direction compared to its dendritic ramifications (like pyramidal cells) (Figure 4.A).

### Phase histograms

The phase histogram counts the phase of tACS at the time when spiking occurs. We computed the tACS phase angle during spike occurrences for each neuron and plotted them for different electric field strengths. In the absence of an electric field, neurons do not fire at a preferred phase with respect to a hypothetical 10Hz oscillation and should thus follow a uniform phase distribution. If tACS synchronizes the firing activity, one should observe a preferred direction in the phase histogram.

### Interspike interval (ISI) histogram

The ISI indicates the time difference distribution between two consecutives spikes. A spike train can be analyzed by measuring the distribution of intervals *s*_*k*_ between two consecutive spikes and plotting them in a histogram. If the spike train is sufficiently long, the ISI can highlight specific firing pattern that can be affected by tACS.

### Data simulation and analysis

Data analysis was performed in MATLAB 2019b (Mathworks, Natick, MA). Neuron modeling and tACS simulation were implemented in the NEURON environment v7.4 (Carnevale & M.L. Hines, 1997).

### High Performance Computing

Analysis was performed on a desktop workstation (Intel i7 9700k, 32 GB RAM, Nvidia RTX 2080). The Neuron simulation workflow was executed on the supercomputing cluster of the University of Minnesota. In total, for each of the 25 neuron morphologies we performed 26 simulations representing a total of 650 simulations. Total computation time was approximately 50,000 CPU-hours.

## Results

### Cortical cell membrane polarization

The response of cortical neurons to tACS depends on the specific cell type. We thus first determined the amount of membrane polarization with respect to the applied electric field strength for each modeled neuron. For this, we used neurons with only passive properties and no synaptic input. We found a linear relationship (mean *R*^2^ = 0.9818 over the set of all neurons) between the soma polarization and the applied electric field strength (see Figure 2A). This linear relationship is fully expected in a low intensity regime due to Ohm’s law.

**Figure 2.**
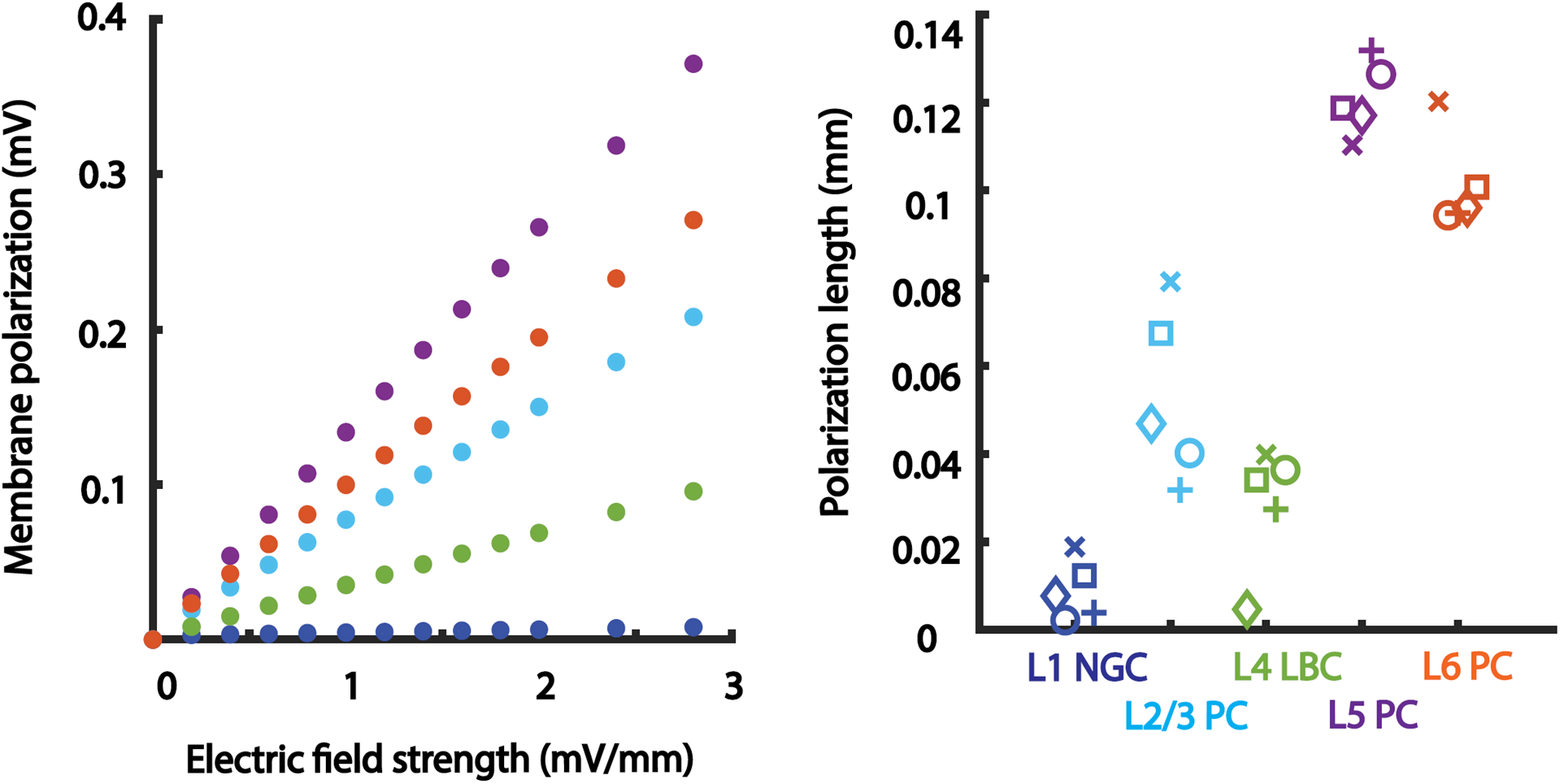
Characterization of subthreshold activity and cortical cell type polarization sensitivity. **(A)** Somatic polarization per applied electric field strength per cell type (one neuron shown per cell type). The slope of each curve determines the subthreshold polarization sensitivity for each neuron. L1 NGC (dark blue) = 0.08 mm, L2/3 PC (light blue) = 0.042 mm, L4 LBC (green) = 0.039 mm, L5 PC (purple) = 0.131 mm, L6 PC (orange) = 0.096 mm. **(B)** The polarization length *λ_p_* (mm) shown for all investigated neurons. The polarization length is an indicator of polarization per unit subthreshold electric field applied.

We then computed the polarization length as the slope of the regression line between soma polarization and electric field strength for all cells (Figure 2B). As expected, non-pyramidal cells had the lowest polarization lengths: L1 NGC (range 0.0077 mm - 0.0189 mm), L4 LBC (range 0.0047 mm - 0.0399 mm). L5 PC show the highest values (range 0.1103 mm – 0.1319 mm) followed by L6 PC (range 0.0944 mm – 0.1203 mm) and L2/3 PC (range 0.0318 mm – 0.0793 mm). We found a significant difference in the polarization length *λ_p_* for interneurons across layers and L5 pyramidal cells (t(13)=-14.72, p=1.72*10^−09^), for interneurons across layers and L6 pyramidal cells (t(13)=-11.30, p=4.25*10^−08^), and between interneurons across layers and L2/3 pyramidal cells (t(13)=-3.73, p=0.0025).

### Effect of tACS on the firing rate

To investigate how the rhythmic membrane de- and hyperpolarization affects spiking activity, we first determined the effect of tACS on the firing rate. For this we compared the firing rate during tACS to the firing rate without tACS. This was done by calculating the ratio of firing rate durinig tACS relative to no-tACS baseline. A value of 1 indicates no change and values smaller or larger than 1 respectively indicate a decrease increase in the firing rate during tACS. We investigated how the firing rate is affected by increasing electric field strength for various cell types. We found that in a low electric field strength regime (< 1 mV/mm), the firing rate changed less than 1% irrespective of the cell type (Figure 3A; see also Supplementary Figure 1 + Table1). For the higher electric field intensities, L2/3 and L5 pyramidal cells show increased firing rate of up to 5%.

**Figure 3.**
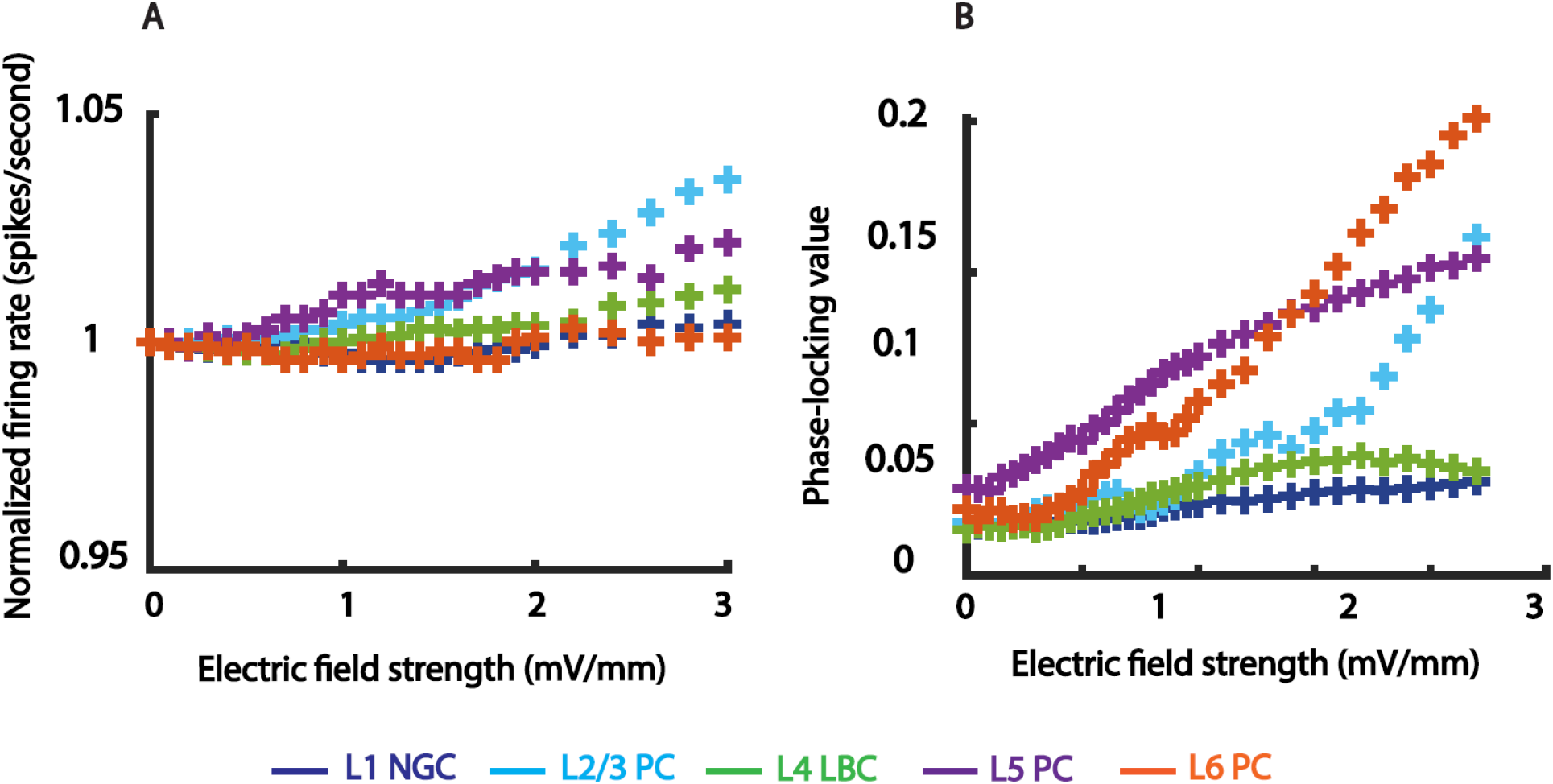
tACS effects on firing rate and phase locking-value (PLV). **A)** Normalized firing rate (change compared to baseline) as a function of electric field strength. Firing rates of different neurons (different colors indicate different cells) are not affected by weak electric fields. Only for high intensities an increase in up to 5% firing rate can occur. **B)** Synchronization of neural firing as measured through the PLV increases for all cells with increasing field strength. Pyramidal cells show a larger increase in PLV than other cell types. This increase is present even in the low electric field strength regime.

**Figure 4.**
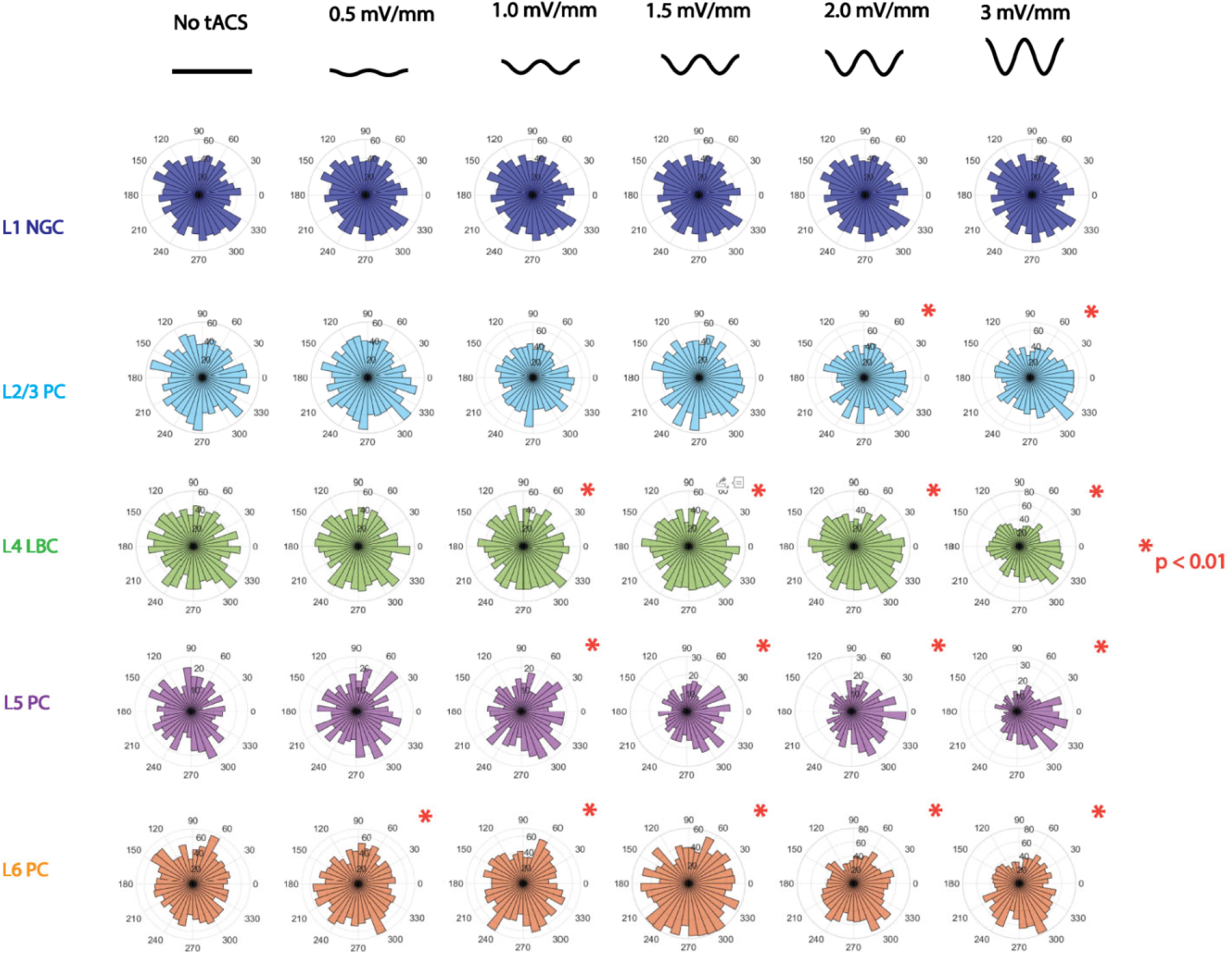
Polar histograms of tACS phase during spike onset. Shown are the phase distributions of spiking with respect to the tACS phase for varying electric field strengths. 0-degrees correspond to the peak of the oscillation. For all cells, tACS affects the timing of spikes with respect to the applied waveform. Phase histograms that differ significantly from a uniform distribution are indicated with a red asterisk (Rayleigh test, p < 0.01). Pyramidal neurons show stronger effects compared to other neurons.

### Effect of tACS on neural entrainment

To quantify the effect tACS has on neural entrainment, we calculated the PLV for all cells as a function of electric field strength. We found that the neural synchronization increases with electric field strength (Figure 1, Figure 3B, see also Supplementary Figure 2 for all cells). Pyramidal neurons exhibit strongest entrainment while the PLV of Layer 1 and Layer 4 neurons increased only slightly (mean slope of 0.0047 and 0.1270 mm/mV respectively while L2/3 PC, L5 PC and L6 PC exhibit a slope of 0.1660, 0.1904 and 0.1139 respectively).

To further investigate the preferential timing of neural firing with respect to the tACS oscillation, we computed phase histograms (Figure 4). We found that with increasing electric field strength, phase histograms deviated more from uniformity and neurons fired closer to the tACS peak. Phase distributions differed from circular uniformity (see Rayleigh test statistics for all neurons in Supplementary Table 2) for electric field strengths as low as 0.5 mV/mm. For an electric field strength of 1mV/mm, only two out of ten non-pyramidal cells exhibit a significant phase preference while seven out of fifteen pyramidal cells exhibit one. The preferred phase angle differed between the investigated neurons. For example, the preferred phase angle for the Layer 6 pyramidal neuron was 300-330 degrees which corresponds to the moment just before the tACS peak. This is in line with experimental literature (Johnson et al., 2020; Krause et al., 2019) showing that preferred phase does not necessarily have to occur at 0-degree (maximum electric field strength) but can also occur at other phase angles.

### Influence of neuron morphology on tACS effects

The neuron morphology is a key feature to understand its response to tACS. In order to quantify the morphology for each neuron, we computed the effective lengths *L_X_* and *L_Y_* as well as their ratio *ρ*_*YX*_ (*L_Y_* divided by *L_X_*). We then investigated the relation between the PLV and the effective length *L_Y_*, and *ρ_YX_* for all 25 neurons at electric field strength of 1mV/mm. We found a correlation coefficient of r(23) = 0.3902 and r(23) = 0.6836 for *L_Y_* and *ρ_YX_* with PLV (see Figure 4B) respectively. Removing the data point with high *ρ_YX_* and PLV still results in a correlation coefficient of r(22) = 0. 4042. This means that a cell that is elongated in the direction of the electric field (*i.e* a large effective length *L_Y_* and a high ratio *ρ_YX_*) will exhibit a large net polarization and thus larger effects on its spiking activity (Radman et al., 2009). The effective length *L_Y_* and the ratio *ρ_YX_* are thus easy-to-derive morphological features that can explain the strong sensitivity of large pyramidal cells to tACS.

### Effects of tACS on interspike intervals

While entrainment is one key mechanism of tACS, it is one specific case of a broader class of changes in spike timing. To investigate changes in spike timing due to tACS in further detail, we computed interspike interval histograms (ISIs) and compared them between tACS on and off conditions.

We focused on the ISI histograms for L5 pyramidal cells showing the strongest entrainment effects (Figure 5). We found that the ISI histograms did not differ significantly between each other (Kruskal-Wallis test, degrees of freedom = 2, p > 0.01). However, we observed that the mean spike timing interval decreased for all cells with increasing field strength (Figure 6). On average, a decrease of 15 milliseconds has been observed from the baseline to 2mV/mm tACS over all neurons (minimum of 4.64ms, maximum of 24.76ms).

**Figure 5.**
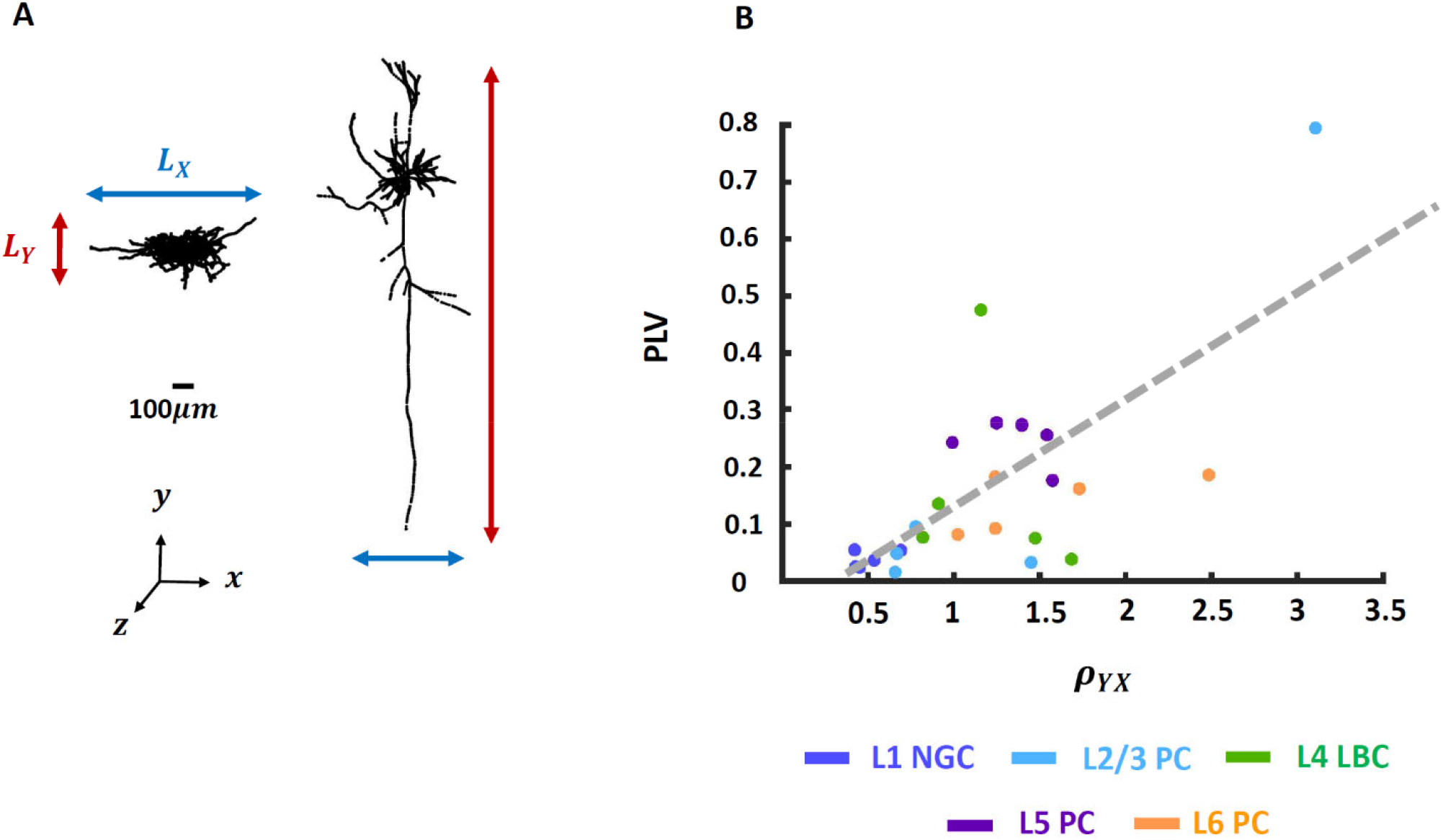
Response to tACS depends on the neuron morphology. **A)** Neurons with a pronounced somato-dendritic axis exhibit a large ratio *ρ_YX_* (*right,* cell from Layer 2/3) while non-pyramidal cells have a low ratio *ρ_YX_* (*left,* cell from Layer 1). **B)** Neurons with higher ratios *ρ_YX_* are more likely to be entrained by tACS as indicated by higher PLV values. Each color represents a specific cell type. PLV values were computed for a field strength of 1 mV/mm.

**Figure 6.**
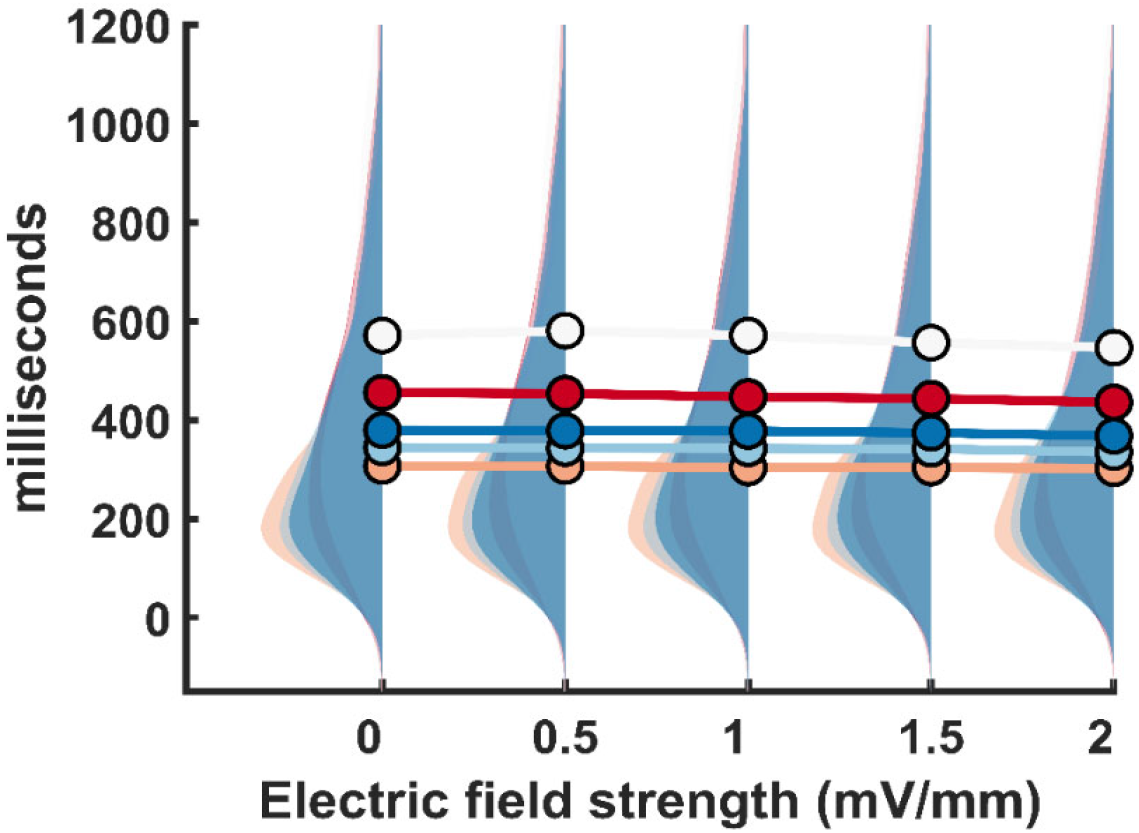
Effects of tACS on inter-spike-interval (ISI) of L5 pyramidal cells. ISI histograms and their mean (solid dots) are shown for different cells (each color indicates a different neuron). With increasing electric field strength, a slight shift towards shorter interspike intervals occurs. Each color line represents the evolution of the mean ISI for each electric field strength.

## Discussion

In this study we developed computational models of morphologically realistic neocortical neurons with ongoing spiking activity to study the effects of tACS electric fields on neural activity. We demonstrate 1) a linear relationship between membrane cell polarization and the applied subthreshold electric field strength, 2) varying polarization lengths across different cell types resulting from their different morphologies 3) no changes in the firing rate of investigated neurons for electric field strengths applicable to human experiments, 4) spike timing changes and neural entrainment occurs at low electric field strengths with largest effects observed in pyramidal neurons.

Our results are in line with previous experimental and computational work highlighting the importance of morphologic features of neurons on somatic membrane polarizations to electric field stimulation (Berzhanskaya et al., 2013; Chan, Hounsgaard, 1988; Radman et al., 2009; Yi et al., 2017). Pyramidal neurons are characterized through asymmetrically placed somas with respect to their dendritic ramifications. In contrast most interneurons have a dendritic tree uniformly distributed in space (Amitai & Connors, 1995). These differences in neuronal morphologies result in differences in polarization across cell types (Bikson et al., 2004; Chan, Hounsgaard, 1988; Francis et al., 2003; Jefferys, 1981) which can be quantified through the polarization length. Interestingly, polarization lengths computed in our study overlap very well with experimentally determined values corroborating the utility of our modeling approach (Radman et al., 2009).

We further investigated the effects of tACS on firing rate and spike timing. We did not observe changes in firing rates for the investigated neurons in an electric field regime applicable to human tACS studies. This is in line with recent in-vivo recordings in non-human primates (Johnson et al., 2020; Krause et al., 2019). Only for electric field strengths > 2 mV/mm, noticeable increases in the firing rate occurred. However, we observed a significant modulation of spike timing across all cell types. Entrainment of spiking was present even for low electric field strengths and increasing for higher amplitudes. Our findings support experimental results in non-human primates showing that electric fields strengths of 0.5 mV/mm are sufficient to cause entrainment of spiking activity (Johnson et al., 2020). These effects were most prominent in pyramidal cells which showed stronger phase-locking of spikes compared to interneurons. The effectiveness of tACS to entrain spiking activity was strongly determined by morphological factors such as the ratio of the effective length of the neuron along the electric field vector to its dendritic arborizations. These findings highlight the importance of taking into account neuron morphology with respect to the applied electric field orientation to optimize tACS physiological outcomes. Our findings suggest that pyramidal neurons are most likely affected by tACS, in particular in an electric field regime present in human studies.

While our model was able to replicate known experimental findings and highlight important mechanisms of tACS, it has some limitations. First, we only chose one excitatory synapse in proximity to the soma (within a 50*μm* range). The synapse location at one proximal dendrite cannot fully take into account nonlinearities and effects arising from dendritic integration (Spruston, 2008). In addition, tACS effects on the single neuron level can be amplified through network effects arising from synchronized synaptic input. Future efforts including multiple sources of excitatory and inhibitory synaptic input in our neuron models could overcome some of these limitations and enable predicting tACS effects occurring in larger neural networks.

In conclusion, our findings advance our mechanistic understanding of tACS effects on single neurons. Our results highlight the importance of neuronal morphology on the susceptibility towards external electric fields. Morphological differences across neurons result in differences in membrane polarization and consequently effects of neuronal entrainment to tACS. Changes in spike timing can already occur at low electric field strengths achievable in human studies, most prominent in pyramidal neurons. In future work, our modeling framework will allow the efficient exploration of tACS parameters thus enabling the optimization of stimulation protocols to maximize their physiological effectiveness.

