## Supplementary_materials for "Effects of transcranial alternating current stimulation on spiking activity in computational models of single neocortical neurons": supplementary_figure1.pdf

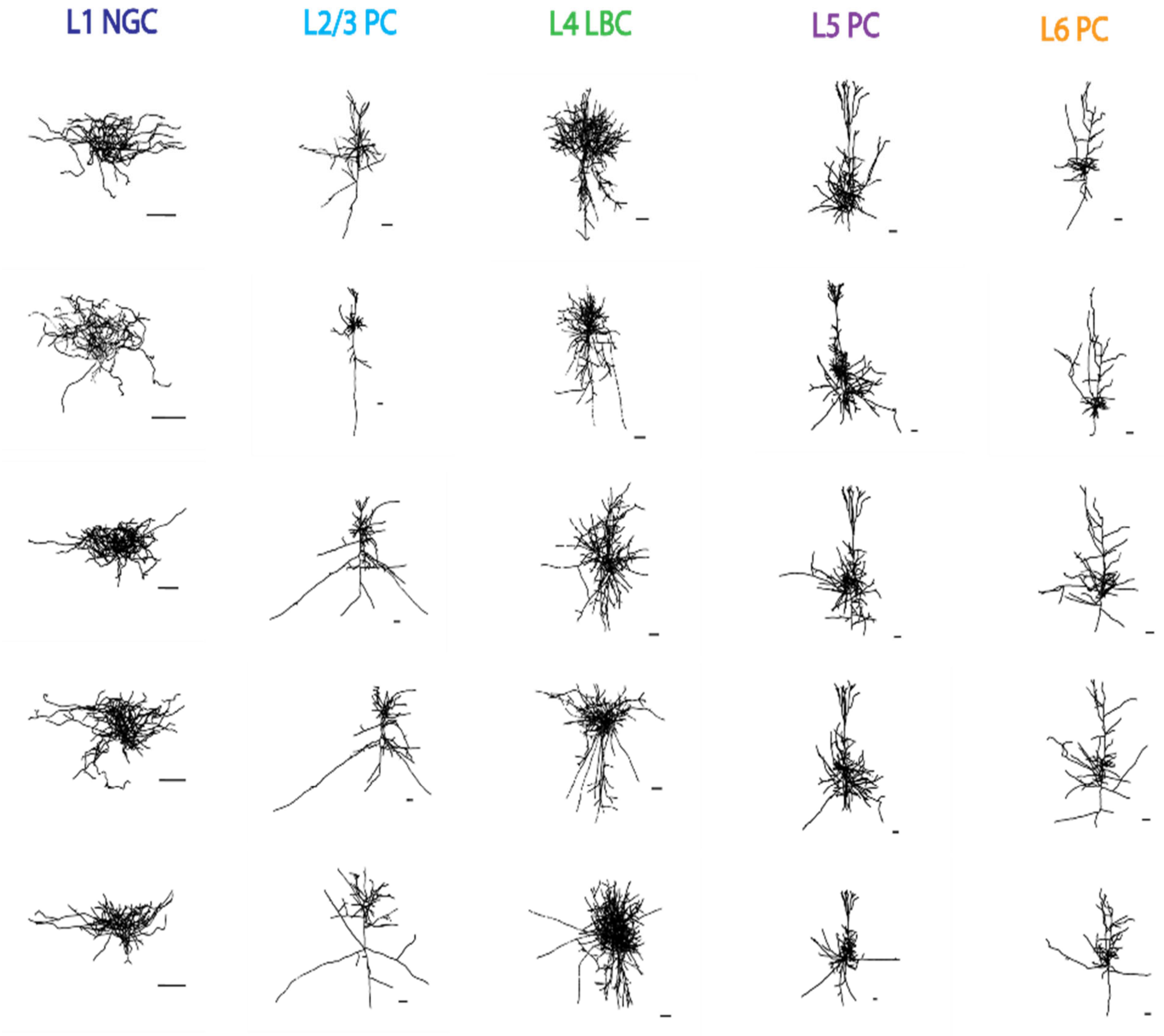


**Supplementary figure 1. Morphologies of neocortical neurons used in this investigation.** Each column displays the five versions of each cell type. Morphological features for each cell type can be observed. Pyramidal cells are larger including long myelinated axon while L1 NGC and L4 LBC cells have larger dendritic ramifications. Scale bars = 100 μm.
