## Supplementary_materials for "Effects of transcranial alternating current stimulation on spiking activity in computational models of single neocortical neurons": supplementary_figure2.pdf

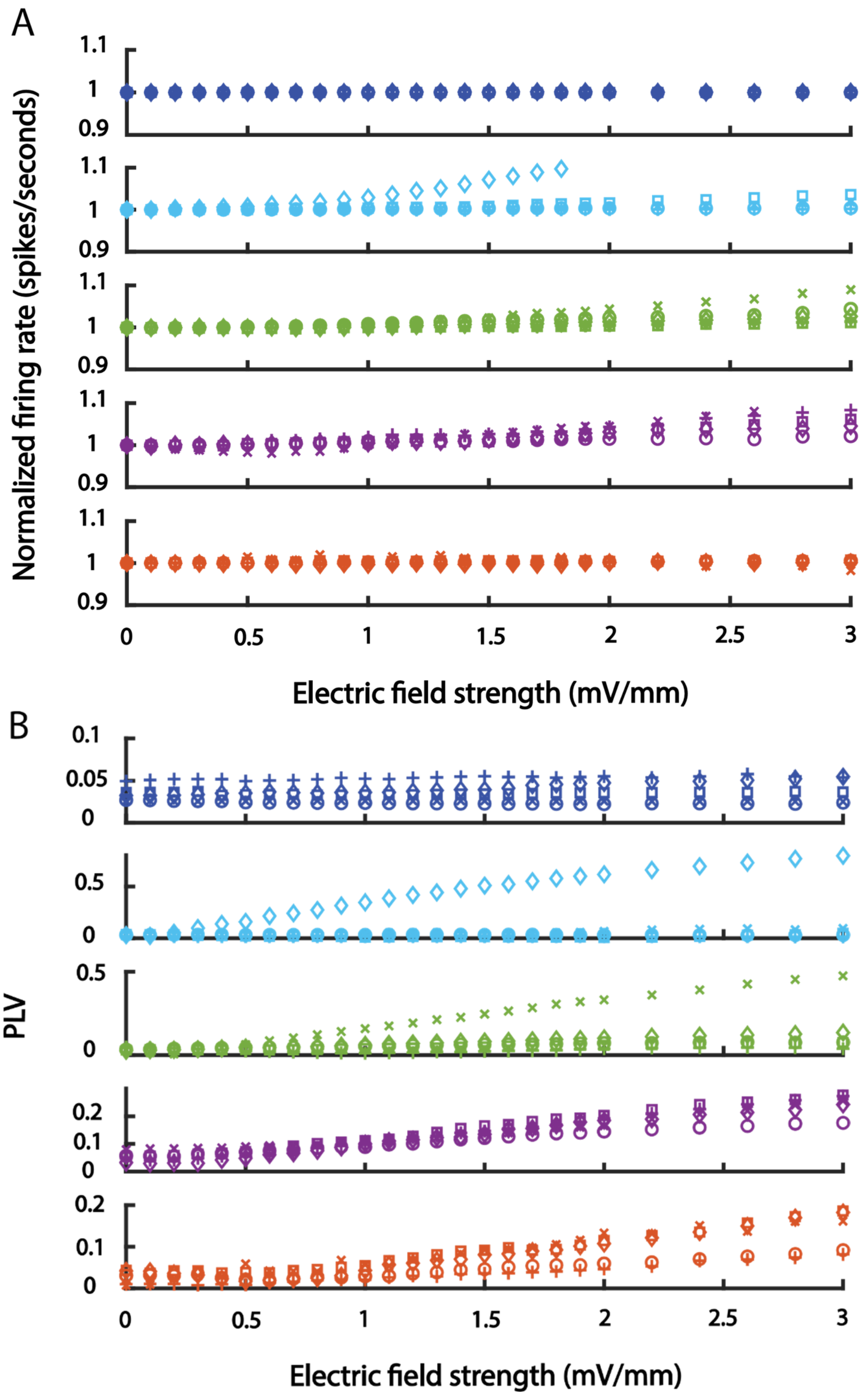


**Supplementary figure 2. tACS effects on firing rate and phase locking-value (PLV) for all cells. (A)**. Normalized firing rate. **(B)**. Phase lag values. Notice the different y-axis scales for the various cell types.
