## Supplementary_materials for "Effects of transcranial alternating current stimulation on spiking activity in computational models of single neocortical neurons": supplementary_table1.pdf

**Supplementary Table 1.** Number of spikes (left) and firing rate (right) during tACS for all 25 cells.

| | Electric field strength (mV/mm) | | | | | | | |
|---|---|---|---|---|---|---|---|---|
| | 0 | | 1 | | 2 | | 3 | |
| | Number of spikes | Firing rate | Number of spikes | Firing rate | Number of spikes | Firing rate | Number of spikes | Firing rate |
| L1 NGC | 1258 | 5.24 | 1255 | 5.23 | 1257 | 5.27 | 1263 | 5.26 |
| | 1913 | 7.97 | 1913 | 7.97 | 1913 | 7.97 | 1913 | 7.97 |
| | 1440 | 6.00 | 1441 | 6.00 | 1441 | 6.00 | 1441 | 6.00 |
| | 1385 | 5.77 | 1385 | 5.77 | 1385 | 5.77 | 1385 | 5.77 |
| | 1746 | 7.28 | 1747 | 7.28 | 1749 | 7.29 | 1748 | 7.28 |
| L2/3 PC | 2963 | 12.35 | 2960 | 12.33 | 2964 | 12.35 | 2968 | 12.37 |
| | 1236 | 5.15 | 1239 | 5.16 | 1240 | 5.17 | 1241 | 5.17 |
| | 1504 | 6.27 | 1505 | 6.27 | 1513 | 6.30 | 1531 | 6.38 |
| | 1514 | 6.31 | 1520 | 6.33 | 1538 | 6.41 | 1568 | 6.76 |
| | 1316 | 5.48 | 1354 | 5.64 | 1466 | 6.11 | 1614 | 6.76 |
| L4 LBC | 1651 | 6.88 | 1654 | 6.89 | 1659 | 6.91 | 1674 | 6.98 |
| | 1340 | 5.58 | 1351 | 5.63 | 1371 | 5.71 | 1399 | 5.83 |
| | 1248 | 5.20 | 1254 | 5.23 | 1303 | 5.43 | 1360 | 5.67 |
| | 1392 | 5.80 | 1392 | 5.80 | 1397 | 5.82 | 1408 | 5.87 |
| | 1472 | 6.13 | 1473 | 6.14 | 1492 | 6.14 | 1511 | 6.30 |
| L5 PC | 527 | 2.20 | 537 | 2.24 | 550 | 2.29 | 571 | 2.38 |
| | 782 | 3.26 | 790 | 3.30 | 794 | 3.31 | 799 | 3.33 |
| | 420 | 1.75 | 420 | 1.75 | 439 | 1.83 | 470 | 1.96 |
| | 696 | 2.90 | 700 | 2.92 | 716 | 2.99 | 739 | 3.08 |
| | 632 | 2.63 | 635 | 2.65 | 652 | 2.78 | 661 | 2.75 |
| L6 PC | 2251 | 9.38 | 2251 | 9.38 | 2252 | 9.38 | 2251 | 9.38 |
| | 1821 | 7.59 | 1825 | 7.60 | 1826 | 7.61 | 1831 | 7.63 |
| | 1657 | 6.90 | 1667 | 6.95 | 1669 | 6.95 | 1627 | 6.78 |
| | 1148 | 4.78 | 1153 | 4.80 | 1152 | 4.80 | 1156 | 4.82 |
| | 1006 | 4.19 | 1002 | 4.18 | 1007 | 4.20 | 1007 | 4.20 |
