## Supplementary_materials for "Effects of transcranial alternating current stimulation on spiking activity in computational models of single neocortical neurons": supplementary_table2.pdf

**Supplementary Table 2.** Statistical details about phase histograms distributions for all cells using a Rayleigh test for various tACS amplitudes. For each cell, the p-value (significative p-value < 0.01) is given by the first line while the z-value is given by the 2nd line.

| | **Amplitude (mV/mm)** | **0** | **0.5** | **1** | **1.5** | **2** | **3** |
|---|---|---|---|---|---|---|---|
| L1 NGC | Cell 1 | 0.0460 | 0.0462 | 0.0318 | 0.0221 | 0.0222 | 0.0257 |
| | | 3.0788 | 3.0733 | 3.4482 | 3.8126 | 3.8059 | 3.6611 |
| | Cell 2 | 0.2498 | 0.3290 | 0.3501 | 0.3799 | 0.3977 | 0.3391 |
| | | 1.3873 | 1.1117 | 1.0495 | 0.9680 | 0.9221 | 1.0817 |
| | Cell 3 | 0.3824 | 0.3902 | 0.3760 | 0.3827 | 0.3893 | 0.4021 |
| | | 0.9615 | 0.9412 | 0.9784 | 0.9606 | 0.9436 | 0.9112 |
| | Cell 4 | 0.1660 | 0.2147 | 0.2193 | 0.1995 | 0.1711 | 0.1597 |
| | | 1.7961 | 1.5386 | 1.5174 | 1.6121 | 1.7655 | 1.8343 |
| | Cell 5 | 0.1628 | 0.1158 | 0.1024 | 0.0686 | 0.0200 | 0.0054 |
| | | 1.8155 | 2.1559 | 2.2792 | 2.6797 | 3.9126 | 5.2251 |
| L2/3 PC | Cell 1 | 0.2866 | 0.4192 | 0.4599 | 0.4575 | 0.4668 | 0.4875 |
| | | 1.2497 | 0.8694 | 0.7767 | 0.7820 | 0.7618 | 0.7184 |
| | Cell 2 | 0.2515 | 0.2330 | 0.2287 | 0.2326 | 0.2703 | 0.2675 |
| | | 1.3802 | 1.4566 | 1.4754 | 1.4582 | 1.3081 | 1.3185 |
| | Cell 3 | 0.1616 | 0.1228 | 0.0328 | 0.0610 | 0.0011 | $8.4106.10^{-7}$ |
| | | 1.8226 | 2.0964 | 3.4162 | 2.7963 | 6.7737 | 13.9612 |
| | Cell 4 | 0.1384 | 0.4418 | 0.8034 | 0.7065 | 0.9023 | 0.0260 |
| | | 1.9773 | 0.8169 | 0.2189 | 0.3475 | 0.1028 | 3.6458 |
| | Cell 5 | 0.2752 | $2.5702.10^{-15}$ | $3.3738.10^{-73}$ | $3.2275.10^{-170}$ | $1.7419.10^{-270}$ | 0 |
| | | 1.2901 | 33.3946 | 161.7927 | 363.4010 | 555.5604 | 1019.0067 |
| L4 LBC | Cell 1 | 0.0244 | 0.4004 | 0.8144 | 0.4406 | 0.0877 | 0.0865 |
| | | 3.7123 | 0.9154 | 0.2053 | 0.8196 | 2.4329 | 2.4463 |
| | Cell 2 | 0.2401 | 0.0727 | 0.0204 | 0.0043 | 0.0008 | 0.0002 |
| | | 1.4265 | 2.6204 | 3.8868 | 5.4429 | 7.1005 | 8.1558 |
| | Cell 3 | 0.3305 | 0.0075 | 0.0075 | $1.3185.10^{-14}$ | $1.3250.10^{-64}$ | $9.1266.10^{-143}$ |

| | | | | | | | |
|---|---|---|---|---|---|---|---|
| | | 1.1070 | 4.8777 | 31.768 | 77.295 | 142.989 | 307.5156 |
| | Cell 4 | 0.2835 | 0.2217 | 0.1240 | 0.0370 | 0.0078 | 0.0003 |
| | | 1.2604 | 1.5065 | 2.0874 | 3.2957 | 4.8439 | 7.9421 |
| | Cell 5 | 0.3804 | 0.0477 | 0.0042 | $5.6785.10^{-5}$ | $2.6219.10^{-7}$ | $8.5983.10^{-13}$ |
| | | 0.9666 | 3.0423 | 5.4557 | 9.7634 | 15.1207 | 27.6635 |
| L5 PC | Cell 1 | 0.2944 | 0.2114 | 0.0031 | $8.2538.10^{-6}$ | $1.2692.10^{-09}$ | $1.5378.10^{-19}$ |
| | | 1.2232 | 1.5542 | 5.7364 | 11.6523 | 20.3127 | 42.5350 |
| | Cell 2 | 0.0758 | 0.0184 | 0.0019 | 0.0019 | $5.8575.10^{-8}$ | $1.3288.10^{-11}$ |
| | | 2.5789 | 3.9896 | 6.2411 | 11.6856 | 16.5761 | 24.8635 |
| | Cell 3 | 0.0702 | 0.0341 | 0.0032 | $2.2416.10^{-4}$ | $5.7144.10^{-6}$ | $2.6858.1014^{4}$ |
| | | 2.6542 | 3.3739 | 5.7083 | 8.3719 | 12.0032 | 30.7620 |
| | Cell 4 | 0.1306 | 0.0278 | 0.0001 | $4.2115.10^{-9}$ | $1.2582.10^{-13}$ | $9.8153.10^{-26}$ |
| | | 2.0352 | 3.5773 | 8.8463 | 19.1667 | 29.4165 | 56.5004 |
| | Cell 5 | 0.5021 | 0.2208 | 0.0016 | $8.4284.10^{-6}$ | $1.9401.10^{-10}$ | $7.2453.10^{-18}$ |
| | | 0.6891 | 1.5104 | 6.4117 | 11.6397 | 22.1884 | 38.9069 |
| L6 PC | Cell 1 | 0.7939 | 0.8484 | 0.3845 | 0.0528 | 0.0081 | $3.0583.10^{-7}$ |
| | | 0.2307 | 0.1644 | 0.9558 | 2.9395 | 4.8081 | 14.9785 |
| | Cell 2 | 0.1711 | 0.4255 | 0.2123 | 0.0214 | 0.0013 | $1.7619.10^{-7}$ |
| | | 1.7651 | 0.8544 | 1.5497 | 3.8431 | 6.6160 | 15.5229 |
| | Cell 3 | 0.9165 | 0.0027 | 0.0048 | $6.1444.10^{-7}$ | $1.2232.10^{-13}$ | $2.3934.10^{-19}$ |
| | | 0.0871 | 5.8944 | 5.3322 | 14.2761 | 29.6085 | 42.6070 |
| | Cell 4 | 0.1130 | 0.1835 | 0.0321 | $5.7043.10^{-6}$ | $2.4404.10^{-07}$ | $3.1922.10^{-18}$ |
| | | 2.1796 | 1.6953 | 3.4356 | 9.7552 | 15.1822 | 39.9522 |
| | Cell 5 | 0.1667 | 0.7526 | 0.1554 | 0.0014 | $7.9706.10^{-7}$ | $2.1083.10^{-15}$ |
| | | 1.7916 | 0.2843 | 1.8613 | 6.5342 | 11.7113 | 33.5261 |
